# Regimes of Complex Lipid Bilayer Phases Induced by Cholesterol Concentration in MD Simulation

**DOI:** 10.1101/432914

**Authors:** George A. Pantelopulos, John E. Straub

## Abstract

Cholesterol is essential to the formation of phase separated lipid domains in membranes. Lipid domains can exist in different thermodynamic phases depending on the molecular composition, and play significant roles in determining structure and function of membrane proteins. We investigate the role of cholesterol in the structure and dynamics of ternary lipid mixtures displaying phase separation using Molecular Dynamics simulations, employing a physiologically-relevant span of cholesterol concentration. We find that cholesterol can induce formation of three regimes of phase behavior, I) miscible liquid disordered bulk, II) phase separated, domain registered coexistence of liquid disordered and liquid ordered and domains, and III) phase separated, domain-anti-registered coexistence of liquid-disordered and newly-identified nanoscopic gel domains composed of cholesterol threads we name “cholesterolic gel” domains. These findings are validated and discussed in the context of current experimental knowledge, models of cholesterol spatial distributions, and models of ternary lipid mixture phase separation.

## Introduction

Certain mixtures of lipids, sterols, and proteins in lipid bilayers laterally separate to two or more domains of unique composition, divided by macroscopically-distinguishable interfaces.(1) Eukaryotic membrane lipid bilayers can be composed of thousands of unique lipids and proteins, and of the many sterols that can exist in membranes, cholesterol (Chol) is ubiquitous.(2)

Mammalian plasma membranes tend to contain 1:3 to 1:1 ratios of cholesterol to phospholipids,(2, 3) though phospholipid membranes can accommodate approximately 66 mol% Chol(4–7) and this upper limit is approached in plasma membranes of astrocyte cells in Alzheimer’s disease patients.(8, 9)

Mixtures of lipid and Chol have been used to understand the phase behavior of complex lipid membranes since the late 1960s. The main lipid phase transition, from gel (s_o_) to liquid crystalline (L_α_) phase, was first characterized through observation of a large peak in heat capacity at the melting temperature (T_m_) in differential scanning calorimetry (DSC) experiments.(10) From the 1970s to 1990 several works using DSC and nuclear magnetic resonance (NMR) found that mixtures of Chol with phospholipids lowered T_m_.(11) These investigations also built evidence for existence of two polymorphs of the L_α_ phase in binary mixtures of di-C 16:0 PC (DPPC) and Chol, finding there to be a phase of low Chol content and a more ordered high Chol content phase using DSC and nuclear magnetic resonance (NMR). These low and high cholesterol content L_α_ polymorphs have come to be known as liquid disordered (L_d_) and liquid ordered (L_o_) phases, respectively.(12) This work culminated in the seminal paper by Vist and Davis who determined the phase diagram of DPPC:Chol mixtures, describing a pure L_d_ phase from 0–10 mol% Chol, a L_o_+ L_d_ coexistence from 10–22.5 mol% Chol, and pure L_o_ phase beyond 22.5 mol% Chol at physiological temperature.(13) Coexistence of spatially separated L_o_ and L_d_ domains on membrane surfaces are evidenced to be central to protein structure, interaction, and function, as many proteins have different affinities for domains.(14, 15)

Formation of domains of lipids in bilayers has been called “lipid phase separation,” “lipid domain formation,” or “lipid raft formation,” each of which has distinct meaning.(1) In general, these terms are used to describe the binary liquid-liquid phase separation that features coexistence of L_d_ and L_o_ phases in the membrane. Over the past 15 years, many investigations have focused on ternary mixtures of Chol with one high and one low melting temperature (T_m_) lipid species.(16) Multiple points on phase diagrams of lipid bilayer phase separations resulting from mixtures of saturated lipids, unsaturated lipids, and cholesterol have been observed using fluorescence spectroscopy,(17, 18, 27–34, 19–26) x-ray scattering,(20, 35–40) atomic force microscopy (AFM),(18, 37, 41, 42) NMR,(24, 30, 32, 43, 44)interferometric scattering,(45) and Raman spectroscopy,(46, 47) allowing us to achieve a general concept of ternary lipid mixture phase diagrams.

In Figure 1 we briefly summarize the current picture of ternary phase diagrams. At relatively lower T (or higher T_m_) s_o_ is evidenced to exist as a macroscopic phase separated state via fluorescence experiments, AFM, and NMR. s_o_ can disappear at physiological temperatures due to presence of Chol(11, 48–51) or unsaturated lipids,(52–55) which lower the T_m_ of saturated lipids. At high (≳40 mol%) Chol concentrations macroscopic phase separations disappear. Critical fluctuations in domain mixing manifest at one or two points in ternary phase diagrams, depending on whether the immiscible region is open or closed.(27, 30, 56–59) Modern fluorescence,(34, 60) x-ray,(39, 61) and AFM experiments(41, 42) have shown that nanoscopic L_o_ and L_d_ domains coexist outside of the miscibiliy gap around 1:1:1 ratio mixtures, as Chol appears to not truly induce the L_o_ phase in unsaturated lipids. x-ray scattering experiments have revealed that ∼60 nm diameter domains of pure Chol domains can coexist with domains of saturated and unsaturated lipids at these high mol% Chol compositions.(37, 50, 62, 63) Beyond the ∼66 mol% solubility limit of Chol in bilayers,(4–7) Chol forms anhydrous crystals in solution.(50, 64) Additionally, though the main phase transition (L_α_ to s_o_) is first order, phase transitions from L_d_ to L_o_ and L_o_ to Chol domains seems to be continuous.

**Figure 1.**
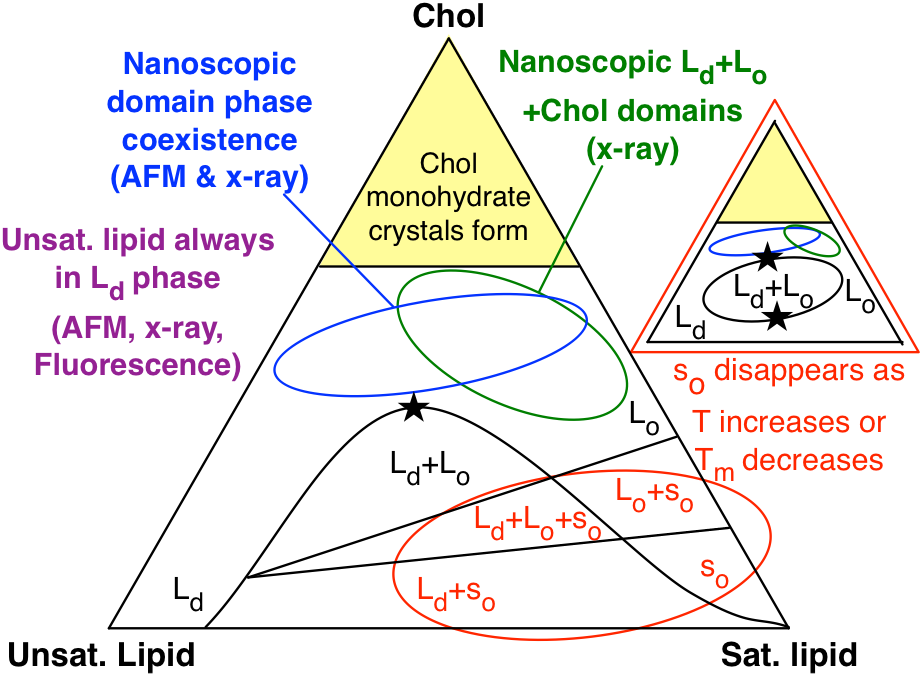
General phase diagram of phase coexistence in ternary lipid bilayers of saturated lipids, unsaturated lipids, and cholesterol (Chol). Liquid disordered (L_d_), liquid ordered (L_o_), gel (s_o_), and pure Chol domains. Black phase boundaries are based on macroscopically-observed phase separations in experiments. s_o_ phase is not present if temperature (T) or main phase transition temperature of saturated lipid (T_m_) are high enough. Regions where nanoscopic lipid domain coexistence have been experimentally observed are noted. Stars indicate where critical mixing is observed in these phase diagrams.

Spatially-resolved, simultaneous knowledge of domains in both membrane leaflets is currently limited. Strong registration of domains between lipid leaflets has been experimentally quantified,(65) however, this registration is only confirmed for macroscopic phase separations of symmetric bilayers, and likely does not occur in mixtures where smaller domains are observed due to complicated surface energies and inter-leaflet interactions.

Many theoretical works have considered the inter-leaflet coupling of lipids and domains, and arguments in favor of inter-leaflet registration or anti-registration of domains have been presented. Preference of domains for local curvature of the membrane surface as well as inter-leaflet interaction between domains can significantly impact the free energy of the membrane. Small, microscopic domains can form that preferably register or anti-register with domains of the opposing leaflet depending on the relative degree of local curvature.(66–73)

Typically, experimental approaches cannot discern the thermodynamic phase and degree of mixing of both lipid leaflets in bilayers of symmetric leaflet composition. Much work has been done to consider inter-leaflet interactions of domains using bilayers of strictly asymmetric leaflet composition, where inter-leaflet domain registration has been observed.(65, 74–76) However, the methods employed, such as use of supported monolayers, may limit the generality of the conclusions of these studies. It is known that Chol strongly prefers to partition to regions of concave curvature.(77) That preferential partitioning may play a role in domain formation and registration.

MD simulations employing the MARTINI coarse grained (CG) model, capable of producing various thermodynamic phases of lipids(78, 79) and lipid phase separation,(80, 81, 90– 92, 82–89) have provided structural insight to inter-leaflet domain interaction. Perlmutter et. al. observed the enhanced local curvature of membranes in the presence of anti-registered domains(90) and Fowler et. al. found that domain registration may occur via a two-step kinetic process of anti-registered domain formation preceding the formation of registered domains.(82) Additionally, Yesylevskyy et. al. demonstrated the preferential partitioning of Chol to regions of locally higher curvature using simulations employing the MARTINI model.(93)

It should be noted that a particular CG lipid may behave differently from its atomistic equivalent.(87) As such, the MARTINI model may be considered as an advanced toy model suitable for understanding general concepts in lipid biophysics, though atomistic models have been shown to reasonably reproduce ternary phase diagrams.(94) With this caveat in mind, CG simulation has contributed to our current understanding of complex membrane phase behavior, though no detailed investigation of the dependence of phase separation and liquid L_d_+L_o_ phase coexistence in ternary mixtures on Chol has been performed.

Here we investigate complex phase separation as a function of Chol concentration in ternary mixtures with DPPC and di-C 18:2 / 18:2 PC (DIPC) lipids, maintained at DPPC:DIPC 1:1 molar ratios, using CG molecular dynamics simulation employing the MARTINI model. Performing simulations at 0, 3, 7, 13, 22, 30, 42, 53, and 61 mol% Chol, we observe that Chol induces three regimes of domain structure at varying concentrations, denoted I) miscible, L_d_ phase, II) domain-registered, macroscopically phase separated L_o_+L_d_ domain coexistence, and III) domain-anti-registered, microscopically phase separated coexistence of L_d_ domains with domains of a newly identified “cholesterolic gel” (s_oc_) phase featuring “threads” of Chol.

Gradual transitions between the three identified phases are observed to be dependent on Chol composition with compositionally unstable mixtures separating regimes sampled at 7 and 42 mol% Chol. The mol% of Chol in DPPC:Chol and DIPC:Chol domains are inferred from the number of lipid-Chol contacts out of all contacts. We find DPPC:Chol domains to rapidly become saturated with Chol, achieving 30% DPPC-Chol contacts by 13 mol% Chol in the membrane, prior to any substantial association of Chol with DIPC. The structure of DPPC, DIPC, and Chol as a function of the percentage of DPPC-Chol and DIPC-Chol contacts is reported in systems at each mol% Chol composition.

This work provides new insight into the role of Chol concentration in complex phase behavior observed and predicted in the past, and provides evidence for a new gel phase that may be thermodynamically stable in membranes of typical Chol concentration.

## Methods

All analysis methods described here were enabled through use of NumPy+SciPy(95, 96) and the MDAnalysis python library,(97, 98) which is built on NumPy+SciPy and Cython.(99) All figures were created using Matplotlib,(100) Omnigraffle, and VMD 1.9.4.(101, 102) All simulations were performed with Gromacs 5.0.4.(103)

### A. MD system construction

The mixture of DPPC, DIPC, and Chol in the MARTINI CG model is able to form macroscopic phase separations, as observed in past investigations of domain formation.(80, 81, 90–92, 82–89) The Chol model of Melo et. al. was used(104) and all other molecules employed the MARTINI v2.0 force field.(105) The insane.py script developed by the Marrink group was used to form three unique initial conditions for 0, 3, 7, 13, 22, 30, 42, 53, and 61 mol% Chol random bilayer mixtures, keeping DPPC and DIPC equimolar.(106) Effectively 38 waters per lipid were used, and 10% of these used MARTINI anti-freeze parameters to prevent spontaneous nucleation of ice droplets. NaCl at 150 mM concentration was used to approximate physiological salt conditions.

We previously found that the MARTINI mixture of DPPC, DIPC, and Chol at 35:35:30 mol% needs to be modelled with more than 1,480 lipids to observe stable macroscopic domain formation.(81) Arguments based on a Flory-Huggins model suggest the need for similarly large systems to observe macroscopic domain formation in other mixtures. In this work, we construct all systems with 3040 lipids, substantially larger than the critical size, such that the PBC will not prevent the observation of domain formation and phase coexistence.

### B. MD Simulation

Each initial configuration was minimized using the Gromacs steepest descent minimizer. Simulation parameters largely corresponded to the “common” parameter set described by DeJong et. al.,(107) and simulations were performed with the same protocol as our previous work.(81) Nonbonded interactions used the Gromacs shifting function applied from 0.9 to 1.2 nm for Lennard-Jones and from 0.0 to 1.2 nm for Coulomb interactions. The velocity rescaling thermostat of Bussi et. al. was used with a coupling time of 1 ps and a reference temperature of 295K.(108) The semi-isotropic Berendsen barostat was used with 1 atm reference pressure, a coupling time of 1 ps, and 3×10^4^ bar^−1^ compressibility, coupling x and y dimensions. The leapfrog integrator was used with a 20 fs time step, employing a “group” neighbor list with a 1.2 nm cutoff, updated every 200 fs. Three replicates of each system were simulated for 11 μs. One set of replicates representing each system condition used San Diego Supercomputer Center (SDSC) Comet resources via the Extreme Science and Engineering Discovery Environment (XSEDE) through startup allocation TG-MCB150142.(109) The other two sets of replicas used the Shared Computing Cluster administered by Boston University’s Research Computing Services.

### C. Atom selections

To analyze the coordinates of lipid head groups in the MARTINI model, we define head groups as the PO4 bead of DPPC and DIPC and the ROH bead of Chol. Lipid tail groups are defined as C2A and C2B for DPPC, D2A and D2B for DIPC, and the centroid of R1, R2, R3, R4, and R5 for Chol.

Chol flip-flops between lipid leaflets were observed in the MARTINI model over the course of simulation. As several of the methods we apply here rely on discrimination between leaflets, we assign Chol to leaflets on a per-frame basis. For each Chol molecule, we find the shortest Chol-lipid head group distance within 1.5 nm, and assign Chol to the leaflet of that lipid.

### D. Order Parameter Analysis

We describe the local concentration of Chol in the membrane in reference to DPPC and DIPC, such that we can infer the content of Chol in lipid domains. We use xy-plane Voronoi tessellations of DPPC, DIPC, and Chol head groups to determine the nearest neighbors, counting the number of DPPC-Chol and DIPC-Chol contacts in the membrane. At equilibrium we infer the Chol composition of L_o_ (L_d_) domains as the percentage of all DPPC-Chol (DIPC-Chol) contacts out of all DPPC-lipid (DIPC-lipid) contacts, noted as <%Chol-DPPC cont.>^eq^ (<%Chol-DIPC cont.>^eq^).

The extent of lateral mixing of DPPC and DIPC in membrane, which is typically used to define phase separation, is described with a binary lateral mixing entropy 
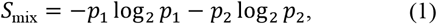
where *p*_1_ is the likelihood of contacts between lipids of the same and *p*_2_ is the likelihood of contacts between lipids of opposite type. This mixing entropy was evaluated by counting the number of DPPC-DPPC, DIPC-DIPC, and DPPC-DIPC head group nearest neighbors determined with Voronoi tessellation (Fig. S1(A)).

Many experimental works infer the transition point between miscible and phase separated states by the inflection point in some order parameter of mixing as a function of concentration or temperature, such as fluorescence intensity. Such inflection points are believed to correspond to point where saturated and unsaturated lipids are 50% miscible. We develop a definition of the 50% miscibility point in 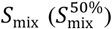 given a binary, periodic 2D system, similar to the Flory-Huggins model used in our previous work.(81) We describe this as a binary mixture in which two pure domains coexist with an ideal mixture that takes up 50% of the system area. The probabilities determining 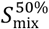 are

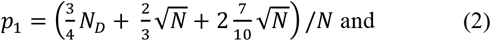

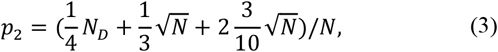
as explained in Supporting Information and Figures S2 and S3. For *N* = 1520, the number of lipids within a leaflet of our simulated systems, we find 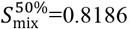.

The registration of lipid domains (Λ) was evaluated by computing the overlapping area of Voronoi tessels centered on lipid tails in each leaflet via Monte Carlo (MC) integration using *N* = 10^5^ points per frame. This was done by evaluating the sum 
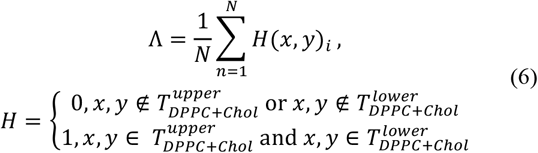
where Λ is counted as number of MC points (x,y) that fall within tessels of DPPC or Chol in the xy-plane of upper and lower leaflets 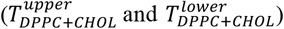 out of the total *N* (Fig S1(B)). To compute the domain overlap describing a random mixture given the same coordinates, we randomly shuffle the chemcial identity of lipids in both leaflets prior to calculation, finding Λ_random_.

The order of lipid tails parallel to the membrane normal was evaluated using the liquid crystal order parameter (*P_2_*) of each lipid tail 
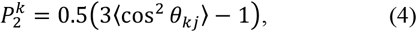
where *k* is the index of the vector representing a lipid tail, *j* is the index of the director vector, and θ_κ*j*_ is the angle between them.

Because the membrane becomes substantially undulated at higher Chol concentrations, using the z-axis as the director vector would not be informative as the structural order of lipid tails is correlated with the membrane surface undulation. In measuring order parameters, we use a plane of best fit for each *k*^th^ lipid tail with its six nearest neighbors (indexed by *l*), found using the singular value decomposition of these coordinates (Fig. S1(C)). *P_2_^k^* is measured using the vector from the GL1 (GL2) to C4A (C4B) beads and the normal vector of this plane of best fit for the *k*^th^ lipid (Figure S1(D)). Here we investigate the average (*P*_2_) for some selection of a number of lipid tails. Only DPPC and DIPC were considered for this analysis as Chol is rigid.

To quantify the order of lipids parallel to the membrane surface, we used the six-fold Nelson-Halperin 2D bond-orientational order parameter 
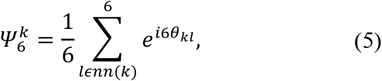
measured using the projections of the *k*^th^ lipid tail and its six nearest neighbor lipid tails (*l ∊ nn* (*k*)) to its plane of best fit to measure θ_*kl*_, the angle between the vector from the *k*^th^ to the *l*^th^ and an arbitrary reference vector, chosen as the projection of the positive x-axis to the plane of best fit of the *k*^th^ lipid tail.

The absolute value of this order parameter measures how well-packed a lipid tail is with its nearest neighbors, with a maximum at 1 (ideal packing). The complex vector of this order parameter describes the bond-orientation of each lipid tail, showing how an ideal hexagon centered on a tail is oriented (Figure S1(E)). We use the angle of the complex vector measured against the reference vector to describe the orientation of each lipid tail in terms of 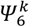 (degrees). We use the average of the absolute value (|Ψ_6_|) for some selection of a number of lipid tails of which DPPC, DIPC, and Chol were considered.

### E. Transleaflet clustering

We evaluated the sizes of domains of DPPC and Chol lipid tails by both number of intra- (*n*) and inter-leaflet (*m*) lipid tails in the domain, using the same hierarchical clustering technique employed in our past work.(81) Inter-leaflet spanning domains of DPPC and Chol were identified by first using hierarchical distance-based clustering of DPPC and Chol tails in one leaflet using a 5.8 Å distance cutoff, assigning the *n* lipid tails discovered using this clustering to the domain. Following this, DPPC and Chol in the opposing leaflet were assigned to the inter-leaflet part of an this domain if the C4A or C4B (DPPC) or C2 (Chol) beads were within 7.0 Å of the tail of any of the *n* lipid tails discovered in the domain. As such, we describe the size of each domain by the number of *n* lipids in one leaflet of the domain, and the *m* lipids in the other leaflet of the domain which were discovered to be in contact with the *n* lipids.

## Results and Discussion

### A. Spatial and structural equilibration

The impact of Chol concentration on the structure and dynamics associated with liquid phase behavior in lipid bilayers was investigated using a CG ternary lipid mixture observed to achieve macroscopic phase separation in molecular dynamics simulation. Simulations of DPPC, DIPC, and Chol lipids at 0, 3, 7, 13, 22, 30, 42, 53, and 61 mol% Chol, maintaining DPPC and DIPC at equimolar ratios, were performed. Three 11 μs replicate trajectories of each system were sampled representing a total of 3 × 9 × 11 μs = 297 μs of simulation. By evaluation of four order parameters, *S*_mix_, Λ, *P*_2_, and |Ψ_6_| (see Methods), we find most systems reach a stationary state by 3 μs. We hereafter refer to the time scale past 6 μs as equilibrium (Figure 2).

**Figure 2.**
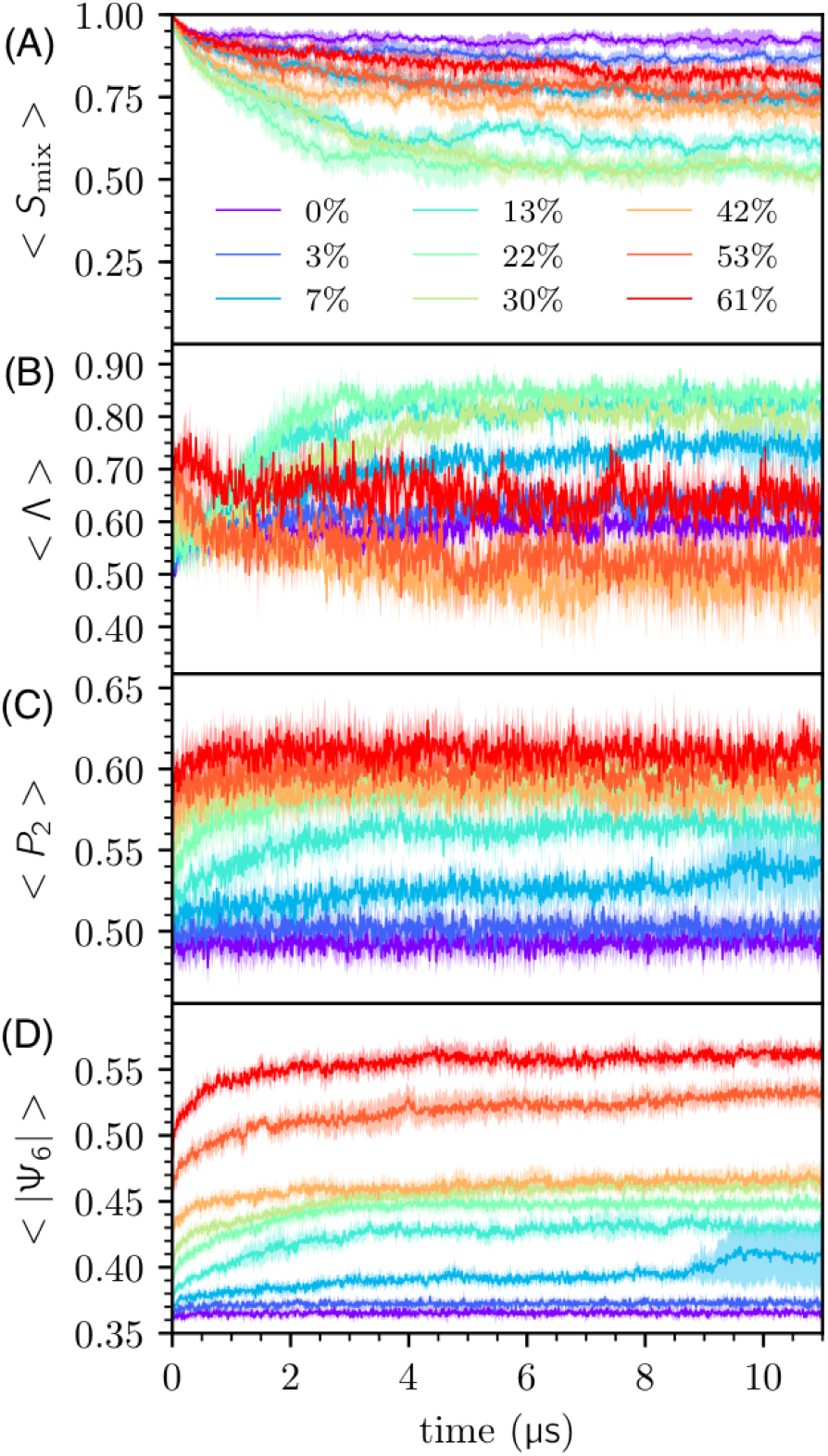
Time series averages of (A) mixing entropy, (B) inter-leaflet domain registration, (C) DPPC *P_2_* order parameter, and (D) absolute value of DPPC bond-orientational order parameter in nine mol% Chol systems.

### B. Three regimes of phase behavior

We observe that the membrane becomes demixed at intermediate Chol concentrations (10–40 mol%) and that domains become registered in the phase separated state, which has previously been confirmed experimentally.(65) Additionally, we observe the well known phenomenon that the membrane becomes more ordered as Chol concentration increases.

Structures drawn from the end of each simulation show a general trend of increasing local curvature as the concentration of Chol increases, ranging from a flat surface (0–13 mol%), to a standing wave (13–42 mol%), to a rough surface (42–61 mol%) (see Figure 3). These changes in morphology appear to be directly related to the partitioning of Chol to regions of concave local curvatures.(77, 110) Local Chol concentration is observed to be spatially correlated with antiregistration of lipid domains at high mol% Chol, and this is implied by the high % of DPPC-Chol contacts accompanying the anti-registration of lipid domains (Figure 4(A), 4(B)). The standing wave observed due to phase separation occurs due to the PBC-spanning size of lipid domains in the phase separated state. This this was observed in other system sizes at 30 mol% Chol in past simulation work.(81) These phase separated domain-spanning undulations have been directly observed in experiment.(111)

**Figure 3.**
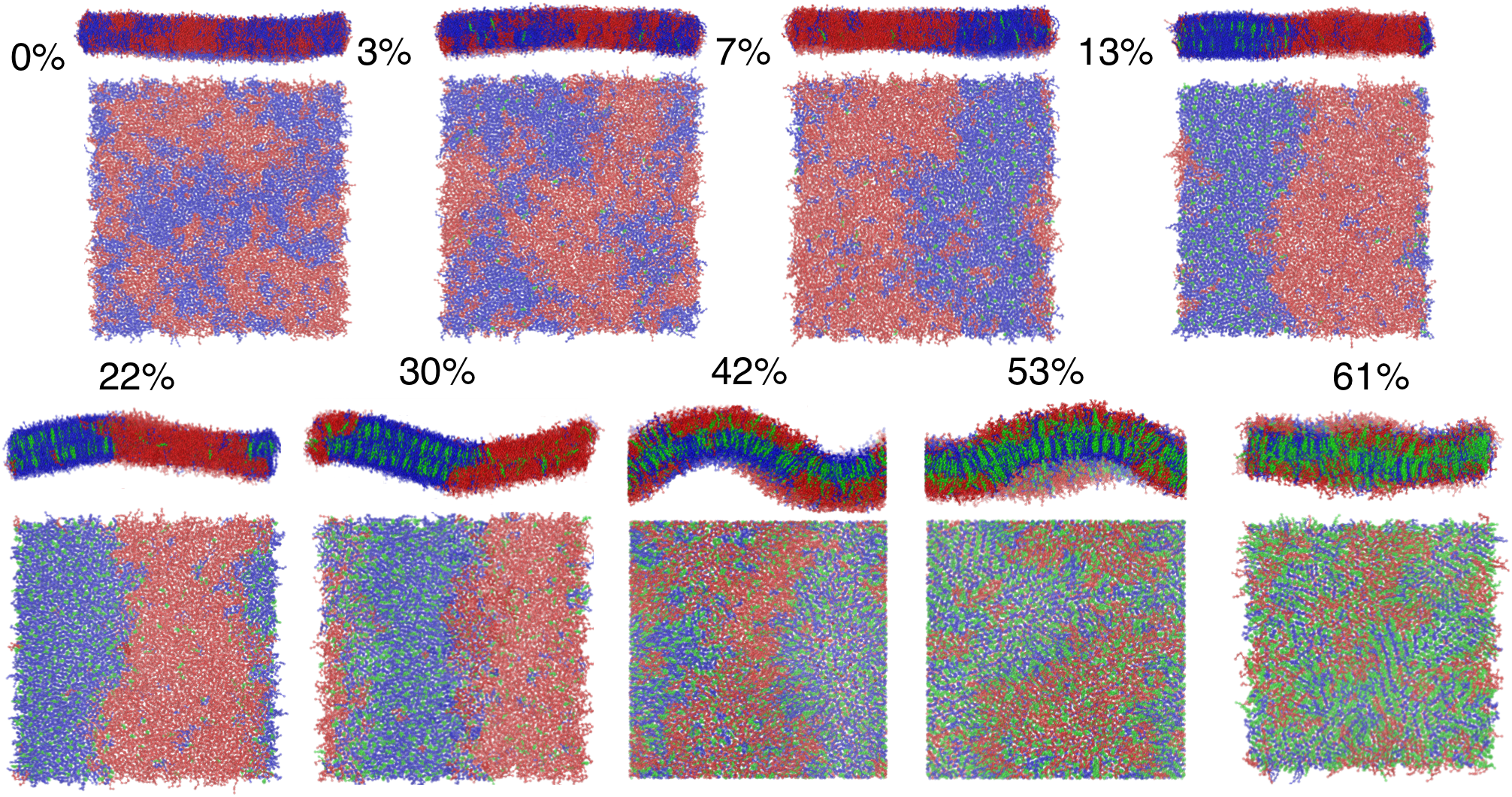
Renderings over the xy-plane and the xz-plane of the final frame of simulation for a trajectory corresponding to each mol% Chol system. DPPC, DIPC, and Chol are rendered blue, red, and green, respectively, using VMD 1.9.4. Bonds are drawn using cg_bonds.tcl from the MARTINI developers.

Chol is observed to preferentially interact with DPPC. Counting the number of DPPC-Chol and DIPC-Chol contacts based on Voronoi tessellations of head groups demonstrate that Chol almost exclusively aggregates with DPPC up to 13 mol% Chol(Figure 4(A)). This colocalization of DPPC and Chol supports L_o_ domain formation, ensuring complete complete formation of L_o_ phase at only 13 mol% Chol, where 30% of all DPPC contacts are made with Chol.(13)Near 20, 33, and 55 mol% Chol the ratio of DPPC-Chol contacts to DIPC-Chol contacts agrees well with recent label-free Raman spectroscopy measurements of Chol partitioning in DPPC:DOPC:Chol monolayers,(46) as well as with x-ray experiments by Chen et al.(40) and Belička et al. at 20 and 24 mol% Chol.(61) The increase of DPPC order parameters as the percentage of DPPC-Chol contacts increase from 0–50% is similar to that observed recently by Wang et. al. (see Figure 3(D), 3(G)).(112) At Chol concentrations surpassing 50% we see that DPPC-Chol contacts exceed the 66% solubility limit of Chol for a bulk membrane.(4–7)

**Figure 4.**
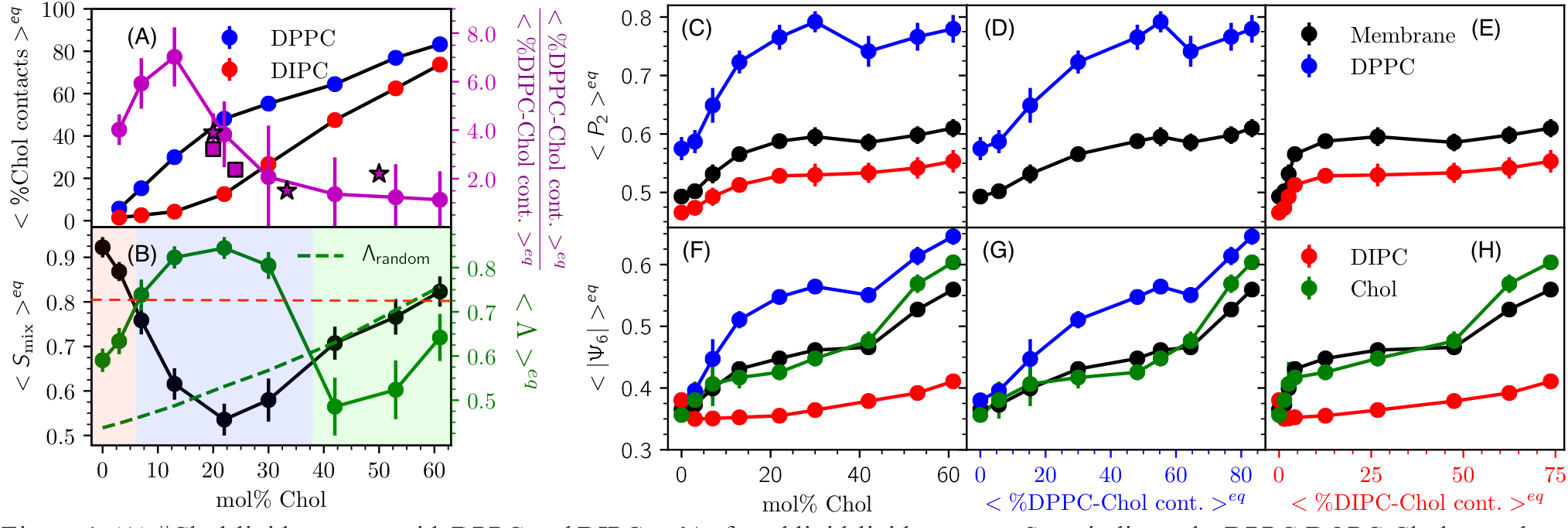
(A) #Chol-lipid contacts with DPPC and DIPC as % of total lipid-lipid contacts. Stars indicate the DPPC:DOPC:Chol monolayer Raman spectra observations of Donaldson and Aguiar.(46) Squares indicate x-ray inferences by Chen et al.(40) and Belička et al.(61) (B) Mixing entropy and inter-leaflet domain registration ratio of %Chol-DPPC to %Chol-DIPC contacts. Shading represents the regimes presenting unique phase behavior labeled I (red), II (blue), and III (green). The dashed red line is the mixing entropy at 50% domain miscibility, 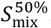. The dashed green line is the equilibrium average domain registration determined from random permutation of chemical identities in these trajectories. Equilibrium DPPC, DIPC, and Chol order parameters as a function of (C and F) mol% Chol (D and G) %Chol-DPPC contacts and (E and H) %Chol-DIPC contacts. All contacts are between head groups.

At high concentrations we observe the formation of anti-registered ordered domains that feature linear aggregates of DPPC and Chol (Figure 3). This structure features repeating face-to-back linear aggregates of Chol separated by a single layer of DPPC tails. This structure is supported by the cholesterol “umbrella model”, in which lipids associate with Chol to prevent solvation of Chol head groups by water. The particular lamellar arrangement of lipids and Chol we observe was previously predicted by umbrella model-inspired lattice simulations developed by Huang and Feigenson.(5) In their simulation model an energetic penalty to Chol-Chol contacts effectively modeled the umbrella effect, and at about 66 mol% threads of Chol were observed to form. These threads minimized the size of Chol aggregates and maintained 2 Chol-Chol contacts. Later, off-lattice simulations by Miao et al. also observed Chol threads.(48) FRET experiments by Parker et al. found that there is a minimum in Chol-Chol contacts about 66 mol% Chol within the range of 45–70 mol%, just below concentrations that would form Chol monohydrate crystals in solution.(6) Parker et al. argued these results support the existence of these Chol thread structures which minimize the cluster size of Chol.

These Chol threads are much like our observed structures above 50 mol% Chol, in which we observe ≳66% of DPPC contacts to be with Chol (Figure 4(A)). We recently performed atomistic simulations to study Chol dimerization structures and found that Chol forms the face-to-back dimers which we observe here with high propensity, suggesting Chol forms threads form not only due to the umbrella effect. (113) Additionally, AFM experiments in ternary mixtures of similar mol% Chol observe the persistence of nanoscopic domains of unknown phase, which may be these domains we report here.(37, 50, 62, 63)

Because these highly ordered domains also exhibit bond-orientational order at the domain length scale (Results section C), making them very similar to the gel phase, we refer this phase as a “cholesterolic gel” (s_oc_)—a lipid gel phase which includes cholesterol. We summarize these observations into three apparent regimes of phase behavior, denoted regime I, miscible L_d_, regime II, domain registered L_d_+L_o_ coexistence, and regime III, domain anti-registered L_d_+s_oc_ coexistence. The transition points between these regimes are discussed in Results section D.

### C. Structure of regimes of phase behavior

At equilibrium the *P*_2_ and |Ψ_6_| order parameters increase up to a “dip” marking the apparent transition between L_o_ to s_oc_ phases at 42 mol% Chol, where 66% of DPPC contacts are shared with Chol (Figure 4(A), 4(C), 4(F)). The s_oc_ phase becomes yet more ordered with higher mol% Chol. The structure of DIPC is generally insensitive to Chol concentrations and the *P*_2_ of DIPC increases only slightly as the L_o_ phase is formed by DPPC and Chol.

The s_oc_ phase is structurally distinct from the L_o_ phase due to the unique lamellar structures formed by DPPC and Chol, manifest in the bond-orientational order of DPPC and Chol. The 3, 22, and 52 mol% Chol systems, representative of regimes I, II, and III, are distinct as characterized by the 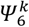 order parameter (Figure 5).

**Figure 5.**
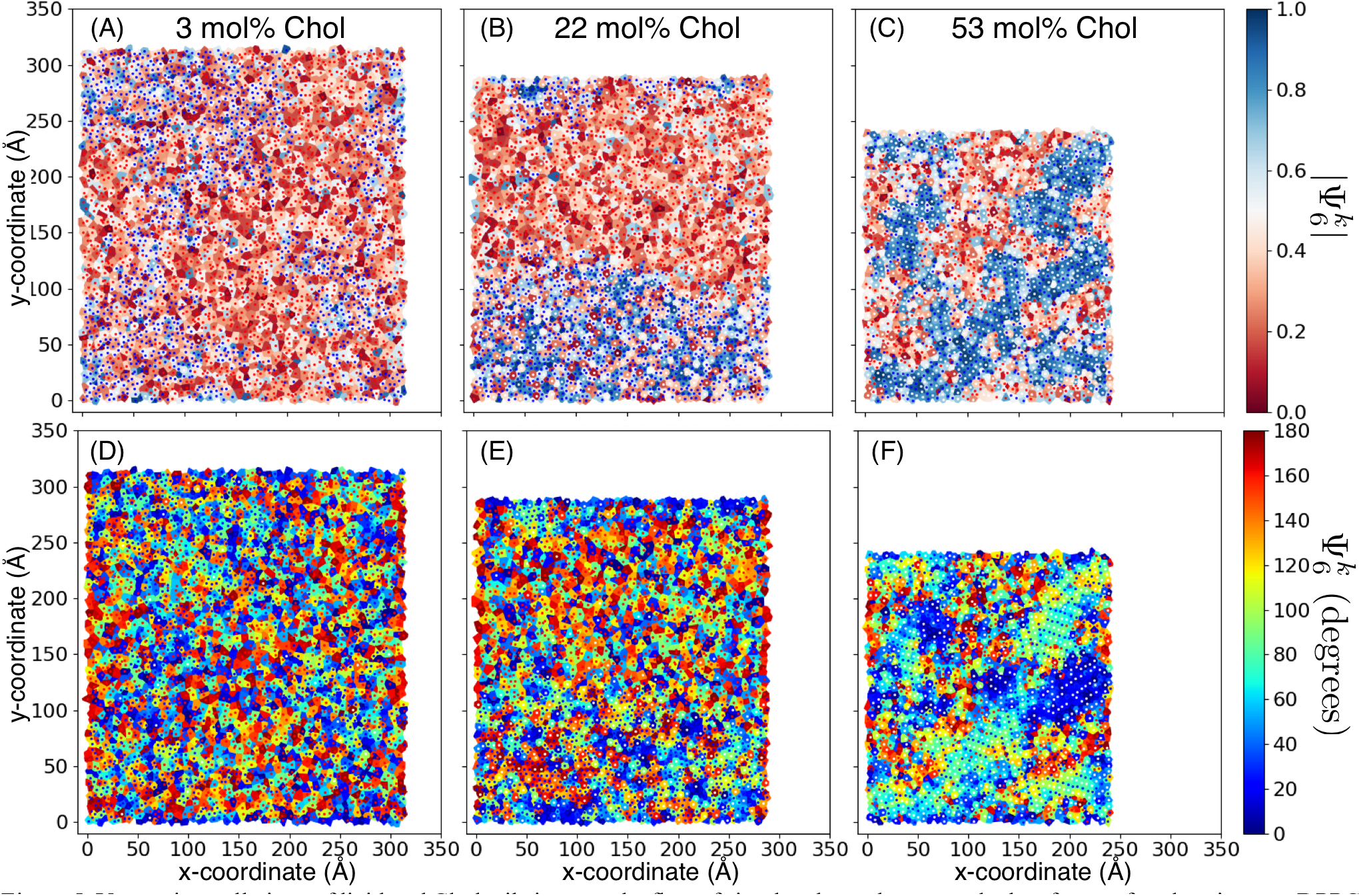
Voronoi tessellations of lipid and Chol tails in upper leaflets of simulated membranes at the last frame of each trajectory. DPPC (blue), DIPC (red), and Chol (white) dots represent tails. Voronoi cells are colored according to the absolute and untransformed value of lipid tail bond-orientational order parameters at (A and D) 3 mol% Chol, (B and E) 22 mol% Chol, and (C and F) 52 mol% Chol.

Inspection of 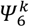 reveals that the L_d_, L_o_, and s_oc_ phases are analogous to the liquid, hexatic, and solid phases of 2D systems. In regime I (0–7 mol% Chol) there is no significant orientational order as measured by 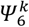 or 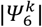 over any length of the system. In regime II (13–42 mol% Chol) 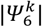 is ordered and correlated over the L_o_ domain. In regime III (53–61 mol% Chol) 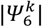 is ordered and 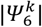 and 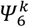 are correlated over the length scale of all s_oc_ domains (Figures S4 an S5).

The s_oc_ phase observed in this study is similar to a gel phase. While gel phases showing “hexatic” order have recently been reported in terms of the Ψ_6_ order parameters,(114, 115) the s_oc_ phase is distinct due to the lamellar arrangement of DPPC and Chol and the effects on the membrane surface that drive domain anti-registration.

Chol, DIPC, and DPPC are evidenced to strongly prefer concave, weakly prefer concave, and weakly prefer convex local curvature on the membrane surface, respectively, as determined via x-ray measurements on monolayers supported by inverted hexagonal phases.(77) Because s_oc_ domains contain Chol in higher concentration, s_oc_ domains may induce concave curvature overall. This would explain the preference for s_oc_ domains to register with L_d_ domains in the opposing leaflet, which are more fluid and contain less Chol. This important difference in domain preferences for local curvature may be accounted for by the theoretical model of Schlomovitz and Schick.(67, 72) However, our results demonstrate that these quantities are sensitive to the concentration of Chol in the membrane, and particularly in each domain, which has not been considered in these models.

### D. Transitions between regimes of phase behavior

In undergoing transitions between phase regimes we expect the system to present fluctuations in local lipid compositions with a disperse distribution of domain sizes. To explore the transitions, we identified transleaflet clusters of DPPC and Chol tails, defining aggregates of *n* intra- and *m* inter-leaflet tails at equilibrium (Figure 6, explanation in Methods, and illustration in Figure S6). We find that the structural order of domains is largely insensitive to domain size (Figure S6(A)), as previously identified in 30 mol% Chol.(81)

**Figure 6.**
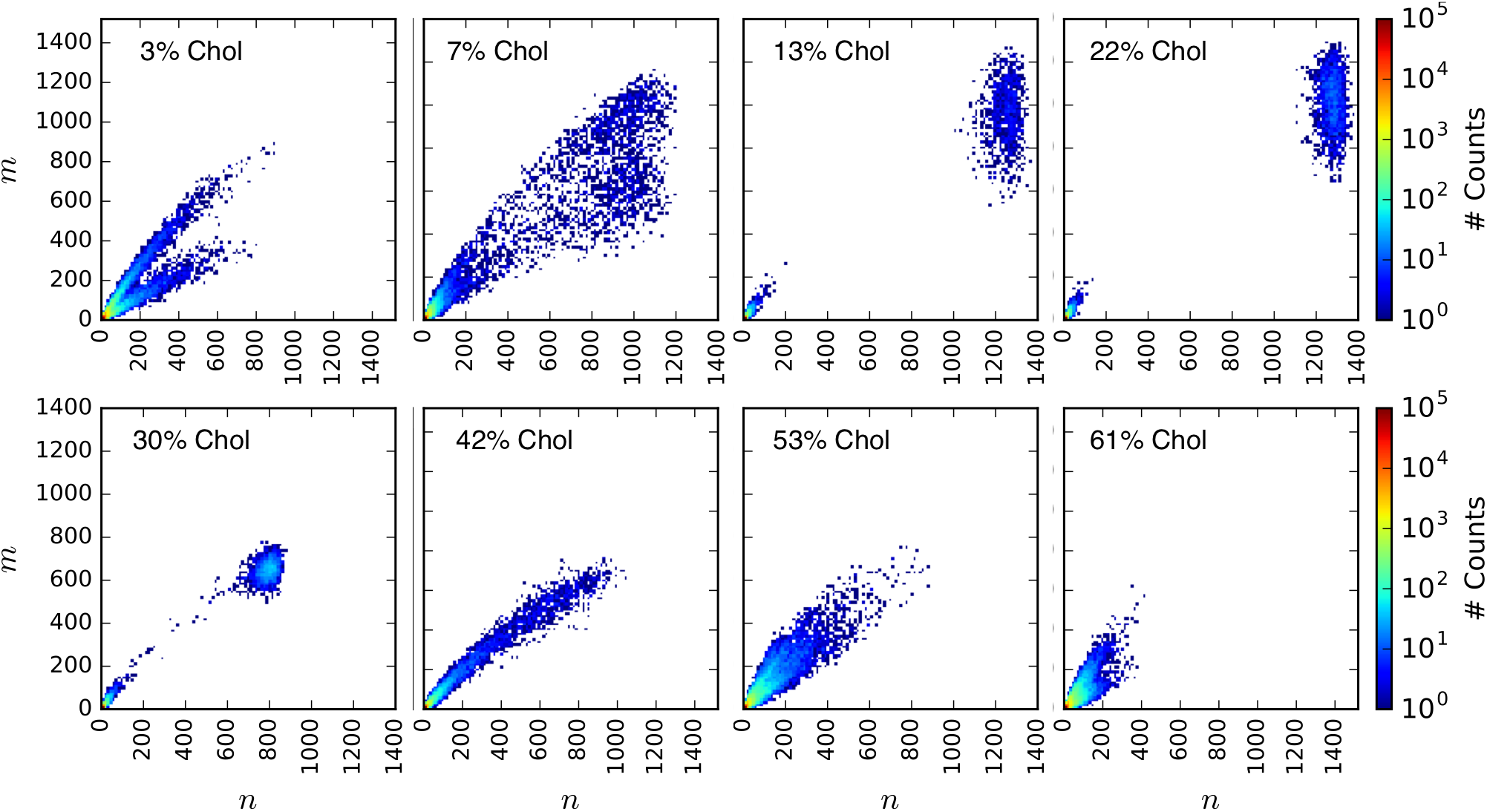
Occurances of intra- (*n*) and inter-leaflet (*m*) lipid tails in transleaflet DPPC-Chol domains at equilibrium in each mol% Chol system.

Examination of transleaflet aggregate sizes in regimes I and III show a bifurcation of slopes of *m(n)*≈2/3 and *m(n)*≈1/3 corresponding to a ∼2/3 domain overlap similar to Figure 4(B). A polydispersity in domain sizes is observed at 7 and 42 mol% Chol. The transition from regimes I to II seems to be well-described by the 50% miscibility point, as the 7 mol% Chol system *S*_mix_, is marginally lower than 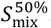 (Figure 4(B)). Additionally, order parameters at 7 mol% Chol show larger fluctuations at equilibrium than other system compositions (Figure 2). The transition observed at 42 mol% Chol is apparently not well-described by 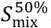. However, the transition of domain overlap, Λ, to below Λ_random_ indicates the onset of anti-registration at 42 mol% Chol(Figure 3(B)), and there is a clear signature of this transition in *P*_2_ and |Ψ_6_| (Figure 4(C-H)). It may be possible to identify the transition to regime III by measurement of domain overlap via a method analogous to our computation of Λ spectroscopically using leaflet-selective deuteration.(74) The transition from nanoscopic domain coexistence to macroscopic phase separation near 7 and vice-versa at 42 mol% Chol approximately fits the DPPC:DOPC:Chol phase diagrams of Veatch et al. and Davis et al., which measure these transitions to occur at approximately 10 and 45 mol% Chol, (33) and 10 and 35 mol% Chol, respectively.(43)

## Conclusions

Via molecular dynamics simulation we have performed a detailed investigation of Chol concentration on phase separation in bilayers formed of ternary lipid mixtures. We observed three regimes of phase behavior, denoted I) miscible L_d_ phase, II) macroscopically phase separated L_d_+L_o_ coexistence featuring registered domains, and III) microscopically phase separated anti-registered domains of L_d_ coexistent with the newly identified liquid cholesterolic gel (s_oc_) phase. These structures were validated by comparison to experimental determinations of Chol partitioning in lipid domains(40, 46, 61) and theoretical expectations of Chol-lipid complex structures at high mol% Chol invoking the umbrella model(5) supported by FRET experiments.(6) We demonstrate the structural difference between these three regimes via order parameters characterizing mixing, domain registration, structural order along the bilayer normal, structural order within the membrane plane, and transleaflet domain sizes. We find regimes I, II and III to manifest distinct differences in bond-orientational order. The s_oc_ phase is found to exhibit 2D bond-orientational order over the length scale of the lipid domains, characterized by lamellar lines of face-to-back threads of Chol and DPPC.

There may be biological implications of the s_oc_ phase for determination of protein structure and function, as proteins can preferentially partition to particular lipid domains. For examples, amyloid precursor processing (APP) is known to change structure due to binding cholesterol(116–119) or changes in membrane thickness,(120–122) which depend on lipid domain composition and structure. APP is processed by α- or β-secretase which residue in different lipid domains.(123–126) If APP is processed by α-secretase, occurring at low mol% Chol, the amyloid cascade will not proceed and production of toxic amyloid beta (associated with Alzhimer’s disease) will not occur. The complex phase behavior induced by cholesterol effects the structure, function, and processing of proteins in Alzheimer’s and other diseases, and will therefore continue to be relevant to our understanding of these disease mechanisms.

These collected observations substantially enhance our understanding of the role of Chol in complex phase behavior in ternary lipid mixtures and provide a framework for exploring structure and dynamics of domain formation in future computational, theoretical, and experimental investigations.

## Associated Content

### Supporting Information

See supporting information for illustrative demonstrations of order parameters *S*_mix_, 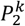, and 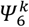, explanation and illustraton of 50% miscibility mapping to *S*_mix_, and visualizations of 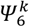 and 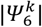 in all systems. Additionally, order parameters measured for transleaflet clusters and illustrative demonstration of the clustering approach used is supplied.

## Author Information

**Author Contributions**

G.A.P. designed the research, performed the research, and analyzed the data. G.A.P. and J.E.S. wrote the manuscript.

## Acknowledgment

G.A.P. thanks the NSF GRFP for support under NSF Grant No. DGE-1247312. J.E.S. acknowledges the generous support of the National Institutes of Health (R01 GM107703). We thank XSEDE, which is supported by National Science Foundation grant number ACI-1548562, and the Shared Computing Cluster of Boston University. We thank Asanga Bandara for assistance with visualization of tessellations and Tetsuro Nagai for interesting conversations regarding the domain miscibility point.

